# Engineered microvessel networks – Longitudinal transcriptomic profiling validates chip-grown tissues as a robust model for studying vasculature

**DOI:** 10.64898/2026.11.05.764625

**Authors:** Luana A. Osório, Ben Hansen, Kinga Suba, Yuyan Yuan, Maja Witowska, Xiang Xu, Md. Noman, Sam Au, Periklis Pantazis, Joseph van Batenburg-Sherwood, Blerina Ahmetaj-Shala, Ildem Akerman, Victoria Salem

## Abstract

Vascularised Organ-on-Chip (OoC) models are promising New Approach Methodologies (NAMs), but their translation requires validation for defined Contexts of Use (CoU). Here, we combine microfluidic microvessel networks (MVNs) with longitudinal bulk and single-cell RNA sequencing to determine how cell provenance influences engineered vascular phenotype. Stromal cell identity produced distinct network architectures and transcriptional trajectories, demonstrating that support cells, commonly treated as interchangeable, represent an important CoU variable. Primary human pancreatic microvascular endothelial cells failed to form stable perfusable networks, but supplementation with 10% Human Umbilical Vein Endothelial Cells (HUVECs) or Blood Outgrowth Endothelial Cells (BOECs) rescued self-assembly. Single-cell transcriptomics identified specialised endothelial populations associated with vascular morphogenesis, including tip, stalk, phalanx, endothelial to mesenchymal transition (EndMT) and pericyte-like endothelial cells. Failed networks lacked coherent endothelial specialisation, whereas rescued networks restored stabilising cell states and revealed FLRT2–UNC5B signalling within pericyte-like HUVECs, recapitulating a pathway implicated in angiogenic development *in vivo*. These findings establish cell provenance as a critical determinant of microvascular NAM performance and demonstrate how transcriptomic validation can define and strengthen the biological relevance of microvessel-on-chip models for specific CoUs.

## Introduction

Endothelial cells (ECs) play an essential role in tissue homeostasis, shaping vascular architecture and regulating function^1–3^. ECs share common characteristics, but display heterogeneity under physiological and pathological conditions, serving highly specialised vascular niches that meet the needs of different organs or processes^4,5^. Replicating tissue-specific functions and morphology in engineered NAMs remains a major research challenge.

The development of organ-specific vasculature involves complex interactions between microenvironmental cues and transcriptional regulation that are not yet well described^4,6,7^. The pancreatic endothelium, for example, specialises according to the various physiological roles of the organ. The endocrine islets of Langerhans possess a dense, fenestrated capillary network that is particularly permeable, facilitating immediate nutrient and hormone exchange required for the tight control of glycaemia. By contrast, the neighbouring pancreatic exocrine parenchyma has lower vessel densities^8,9^. Of growing interest is the bi-directional crosstalk between a given vasculature and its parenchyma that regulates tissue function, and ECs are increasingly considered therapeutic drivers in regenerative medicine^10,11^.

Blood vessel formation and maintenance is a complex process that involves the specialisation of ECs into tip, stalk and phalanx cells. With angiogenic stimuli, tip cells lead the growing vessel through induction of extracellular gradients of vascular endothelial growth factor A (VEGF-A)^12^. Stalk cells follow tip cells and show a high proliferative capacity with the ability to secrete extracellular matrix (ECM) and initiate lumen formation. Role selection for each of these cells is dependent on distinct metabolic and signalling pathways. For example, tip cells have been shown to have higher levels of delta-like ligand 4 (Dll4) when compared to stalk cells which, in turn, upregulate Notch signalling. Phalanx cells on the other hand are quiescent ECs that form the walls of mature vessels but can differentiate into stalk-like cells upon exposure to angiogenic stimuli^12–14^.

There is a growing consensus that during injury, Endothelial Colony Forming Cells (ECFCs) serve as a recruitable reservoir of circulating ECs capable of promoting *in vivo* neovascularisation and tissue repair^15–17^. These endothelial progenitor cells, characterised by the expression of primitive haematopoietic markers (CD133, CD34) as well as established endothelial cell markers (CD31, CDH5, KDR), can differentiate into mature endothelium and initiate vascularisation^18–21^. ECFC transplants have been tested in limb ischaemia to promote neovascularisation^22^ and delivered intravenously to target traumatic brain injury^23^. These cells have also gained traction as models for endothelial pathology because they retain disease-specific transcriptomic features. Blood Outgrowth Endothelial Cells (BOECs) have been described as the *in vitro* expanded and differentiated progeny of circulating ECFCs, which can be isolated non-invasively from the Peripheral Blood Mononuclear Cell (PBMC) fraction of a human venous blood sample^24^. They represent a powerful modelling tool for personalised medicine ^25,26^, for example BOECs grown from patients with type 1 diabetes (T1D) retain a phenotype that serves as a model for diabetic microvasculopathy^27^.

While animal models fail to faithfully recapitulate the complexity of human disease, the development of physiologically-relevant vascularised *in vitro* models is challenged by the difficulty of reproducing the intricate cellular and microenvironmental aspects of human tissues^28–30^. As a result, there has been a growing interest in Organ-on-Chip (OoC) models of human microvasculature to study vasculogenesis, sprouting-tip dynamics, angiogenesis, immune trafficking, tumour intravasation, and microenvironmental crosstalk^14,31–33^.

Current studies using engineered microvessel networks (MVNs) generally rely on the exceptional capability of Human Umbilical Vein Endothelial Cells (HUVECs) to self-assemble *in vitro*. Indeed, these commercially available, well-characterised and highly proliferative endothelial cells have come to serve as the standard for *in vitro* vascular models ^14,34–39^. Aside from using HUVECs, other groups have achieved vascular self-assembly into engineered perfusable networks using mature endothelial cell lines, such as Human Aortic Endothelial Cells (HAECs) ^40^. Pluripotent stem cell derived ECs, on the other hand, have been used to form vascular organoids that are devoid of perfusable lumens^41,42^. These cellular models for vasculogenesis provide valuable insight, but ultimately lack organotypic complexity^1,4,43^.

In self-assembled MVNs, just as in the microvasculature *in vivo,* ECs only make up part of the picture. Of equal importance is their associated structural support cell type. Stromal cells (SCs) are largely responsible for the secretion of ECM, growth factors and cytokines, and physical support for tissue architecture ^6,44,45^. Various SCs, including Dermal Fibroblasts (DFs), Normal Human Lung Fibroblasts (nHLFs), and Mesenchymal Stem Cells (MSCs) of both adipose and bone marrow provenance, have been seeded simultaneously with HUVECs to promote self-assembly on chips ^6,46^. Although there is some evidence that the SC type plays a key role in both regulating engineered vascular morphology and long-term stability^6,38,46^, the specifics of endothelial-stromal crosstalk remain largely unexplored. To advance the field toward more physiologically relevant model systems, it is critical that MVNs derived from these divergent cells are characterised functionally and morphologically. However, previous efforts to map the transcriptomic landscape of engineered MVNs have been constrained by technical challenges related to the extraction of genetic material from lab-grown vasculature.

Here we systematically demonstrate how key subpopulations of ECs or SCs inform overall MVN structure and function. We show that the addition of small proportions of HUVECs or BOECs can rescue the self-assembly properties of terminally differentiated ECs to engineer microvasculatures that are more physiologically-relevant to the native tissue of interest, utilising pseudotemporal transcriptomic exploration of variant vasculature on a chip.

## Results

### MVN architecture in a microfluidic device is dependent on support cell type

Most studies with vascularised OoC platforms use stromal cells interchangeably, with limited justification for the choice. Recent studies have shown that SC identity influences endothelial morphology and ECM production^6,46^. In order to address this knowledge gap, we designed a 3-channel microfluidic device for the generation of MVNs, where EC type, in this case HUVECs, was maintained throughout, and SC type was altered between experiments (Fig. 1a, b), to compare adipose-derived MSCs, nHLFs, and Human Pancreatic Fibroblasts (HPFs) (Fig. 1c). The central channel geometry was constrained in height (200 µm) to encourage MVN formation within a small plane to allow for precise focus on SC-dependent morphological divergence. ECs and SCs embedded within a fibrin hydrogel were allowed 30 minutes of gelation at 37°C, followed by immediate administration of media via hydrostatic pressure gradients. For the first 24 hours, the MVNs grew under static conditions, followed by the subsequent hydrostatic pressure gradient of ∼500 Pa, alternating daily in directionality. All cells were loaded at consistent EC and SC concentrations (reported in Methods section). Vascular self-assembly was evident over the course of the first 24 hours, nascent vessel perfusion was possible after 3 days of culture, and network morphology stabilised after 5 days, at which point the devices were perfused with microbeads and dextran (Fig. 1d).

**Figure 1.**
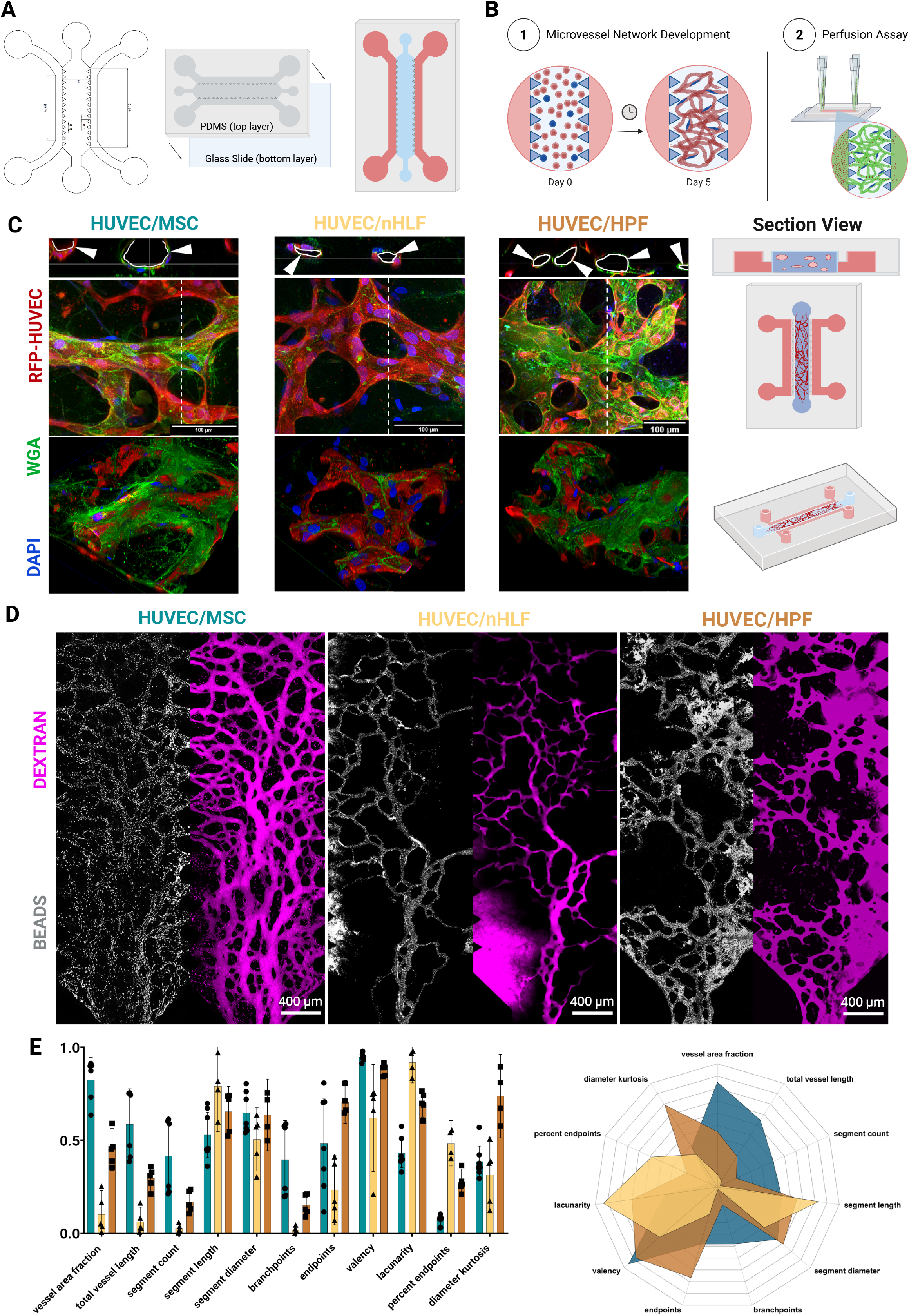
Stromal cell identity regulates microvessel network architecture and perfusability in a microfluidic device. (A) The microfluidic device consists of a gel loading channel (central channel) separated from the adjacent media channels by structural posts, design measurements shown in mm. (B) Temporal schematic of MVN self-assembly over 5 days, followed by perfusion of dextran and microbeads for network quantification analysis. (C) Maximum intensity projection (MIP) of microvessel networks at day 5 with RFP-HUVECs (red), WGA (green) and DAPI (blue) at 63x magnification. White arrowheads indicate representative junctions or lumenised regions. Scale bars 100 μm. Three-dimensional reconstructions of above MIP. Schematic of the section imaged within the microfluidic device. (D) Maximum intensity projection of 70kDa dextran (magenta) and 1 µm microbead (white) perfusion through the vascular networks on day 5 for co-cultures of HUVEC/MSC (n=7), HUVEC/HPF (n=5) and HUVEC/nHLF (n=5). Perfusion is performed unidirectionally from one medium channel to the other over the period of 5 minutes at 10x magnification. Scale Bar, 400 µm. (E) Quantitative network morphology analysis studying vessel area fraction, vessel length, segment count, segment length, segment diameter, branchpoints, endpoints, valency, lacunarity, percent endpoints, and diameter kurtosis between different stromal conditions. All data are presented as means with 95% confidence intervals (CI).

Vessel network analysis (VNA) was performed using bespoke algorithms (process described in Supplementary Figure 1a) that quantitatively defined the morphology of emergent, engineered vasculature (Fig. 1e and Supplementary Figure 1b). Briefly, HUVEC/MSC networks achieved the highest vessel area fraction (57%) and segment count per µm^2^ (8.93x10^-5^) and the lowest endpoint percentage (6%). Further, their average branchpoint count per µm^2^ (8.23x10^-5^) and average segment length of 123 µm were comparable to native vessels ^47,48^, although their vessel diameters (111 µm) were consistently larger than equivalent values reported *in vivo*. In fact, only HUVEC networks grown with nHLFs displayed vessel diameters (86 µm) approaching those found *in vivo*. MVNs grown using HPFs demonstrated the highest average number of endpoints (7.93x10^-6^ per µm^2^), suggesting the presence of blind ends as is visually apparent (Fig. 1d).

### Longitudinal bulk RNA sequencing reveals the transcriptomic drivers of SC dependent MVN morphology

To understand the key drivers of divergent MVN morphology on a transcriptomic level, we performed bulk RNA sequencing at distinct timepoints during MVN morphogenesis. To this end, sequenced tissue extracted from MVNs grown with HUVECs and adipose-derived MSCs (Adipose); HUVECs and nHLFs (Lung) and HUVECs with HPFs (Pancreatic) was analysed over time. Bulk RNA was sequenced on day 0, directly taken from monolayer cell cultures, on day 3 following the formation of nascent vessel networks and on day 5 when MVNs were stable and reproducibly perfusable (Fig. 2a). Principal Component Analysis (PCA) of divergent MVNs netted neat clustering by cell type and time point (Fig. 2b). PCA suggests that for all engineered MVNs, cells converge towards a ‘network’ phenotype, as day 3 clusters are more similar to day 5 than either are to day 0. Gene Ontology (GO) analysis of the growing engineered MVNs revealed distinct transcription pathways (Fig. 2c and Supplementary Figure 2b). All pathways referenced below were identified via GO and corroborating genes extracted subsequently from corresponding pathways.

**Figure 2.**
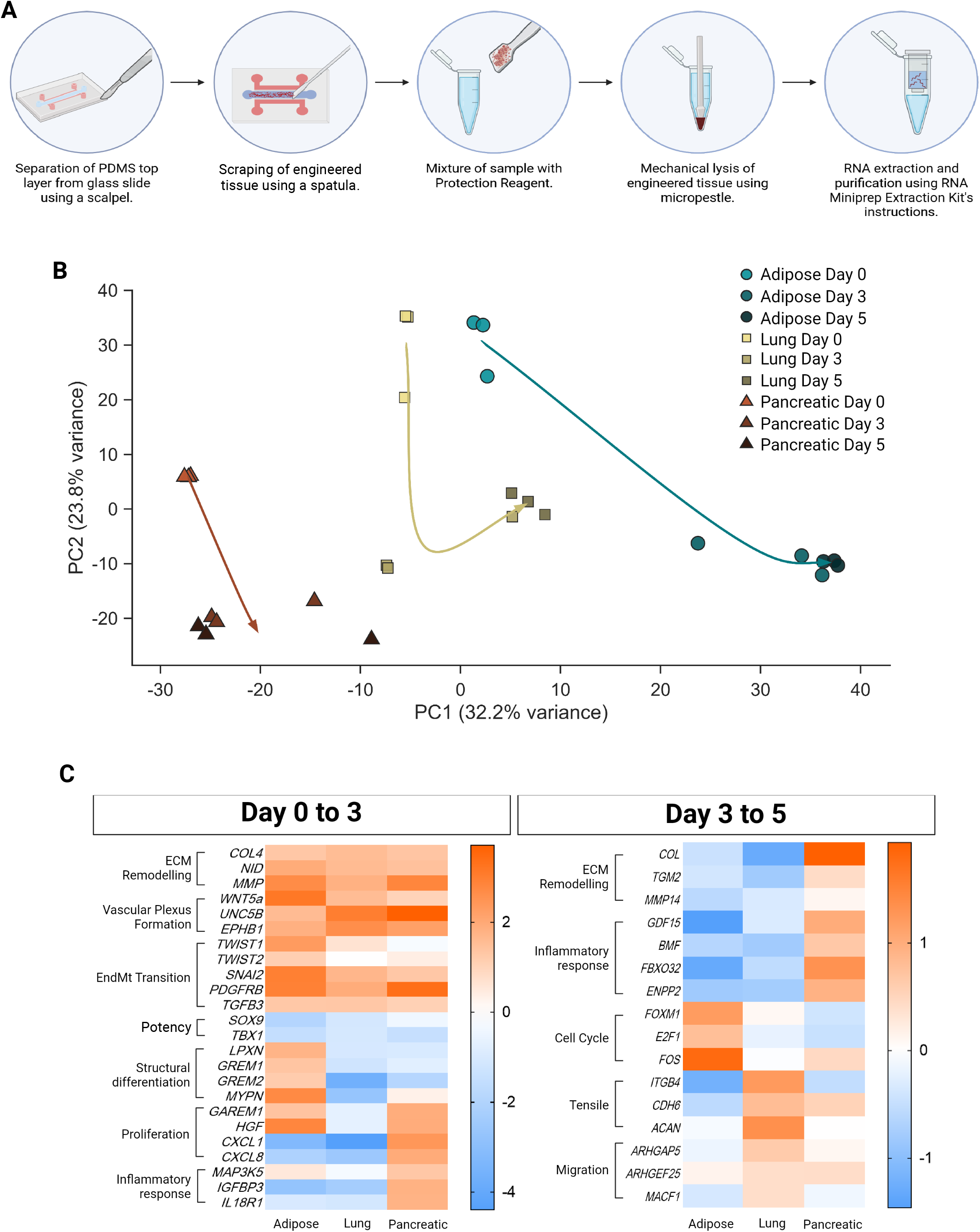
Bulk RNA-sequencing of engineered microvascular tissues reveals tissue-specific and time-dependent gene expression. (A) Schematic overview of sample processing and RNA extraction workflow. (B) Principal component analysis (PCA) of stromal co-cultured tissues, with arrows inferring longitudinal developmental trajectories. (C) Heatmap of differentially expressed genes between day 0-3 and day 3-5 within each tissue condition, scale bars represent log fold change.

From day 0 to day 3, all networks upregulated genes associated with ECM deposition and organisation (COLs, NIDs, MMPs), vascular plexus formation (WNT5A, UNC5B, EPHB1), and cell transition, particularly endothelial-to-mesenchymal transition (TWIST1,2, SNAI2). Genes associated with multipotency (SOX9, TBX1) were downregulated, suggesting that cellular phenotype had become more consolidated. HUVEC/MSC networks in particular supported muscle and mesenchymal cell differentiation (LPXN, GREM1,2, MYPN), whereas HUVEC/nHLF networks downregulated these same pathways, favouring instead proteoglycan biosynthesis, a major ECM component found in the lung^49^. HUVEC/HPF networks, whilst especially metabolically active and proliferative during this period (GAREM1, HGF, CXCL1,8), were even more aggressive in their suppression of genes related to muscle, cartilage, and connective tissue development and differentiation. Broadly, developing vascular networks are known to express pro-inflammatory pathways on GO as inflammation and angiogenesis during wound healing are mutually reinforcing processes. However, HUVEC/HPF networks in particular significantly downregulated nitric oxide (NO) production and regulatory pathways, as well as selectively upregulating the pro-inflammatory genes MAP3K5, IGFBP3 and IL18R1.

From day 3 to day 5, HUVEC/MSC networks suppressed ECM expression (COL, TGM2, MMP14) and inflammatory/senescence pathways (GDF15, BMF, FBXO32, ENPP2), while upregulating genes associated with cell division and cell cycle progression (FOXM1, E2F1, FOS). HUVEC/nHLF networks encouraged cell-cell and cell-substrate adhesion (ITGB4, CDH6, ACAN), highly upregulating migration (ARHGAP5, ARHGEF25, MACF1), while similarly downregulating inflammatory markers. Notably, all networks downregulated ECM pathways except for HUVEC/HPFs, which continued to highly express them. These MVNs continued to express pathways associated with high cell stress, inflammation, and activity.

### MVN self-assembly potential in organotypic primary ECs is rescued by the addition of small sub-populations of proliferative or immature ECs

Terminally differentiated, primary organotypic ECs are difficult to culture and even more challenging to engineer into perfusable vessels ^50,51^. Of particular interest to diabetes researchers are Human Pancreatic Microvascular Endothelial Cells (HPaMECs), isolated from the human pancreas. Attempts to model pancreatic tissue, alongside the appropriate vasculature, are in their infancy but highly sought after as endothelial and β-cell function are intimately linked^52–54^. A more comprehensive understanding of pancreatic vascular biology is therefore indispensable in order to advance T1D research. When HPaMECs were co-cultured with MSCs on microfluidic devices, we found that HPaMECs failed to convincingly and reproducibly organise into lumenised vessels (Fig. 3a,b), despite some instances where network perfusion was partially achieved (Supplementary Figure 3b). Even in instances where MVNs were partially perfusable, the emergent networks were incomplete and disintegrated in multiple areas (n=6). Surprisingly, when HPaMEC and MSC co-cultures were supplemented with an additional small population (10%) of either HUVECs or BOECs, self-assembly was achieved in one hundred percent of cases (n=11). This concentration of supplemental ECs was so low as to be nearly indiscernible on the chips by day 4 (Supplementary Figure 3c). Immunofluorescence staining of HPaMEC/RFP-HUVEC chips on day 5 revealed <5% remnant population by area of RFP-HUVEC, and suggested that a proportion of these cells adopt a ‘pericyte-like’ phenotype where they line the abluminal side of MVNs, rather than their expected inclusion in a chimeric vessel wall. Of particular interest was the finding that differences in network morphology between HUVECs and BOECs were conserved even in these chips (Fig. 3c).

**Figure 3.**
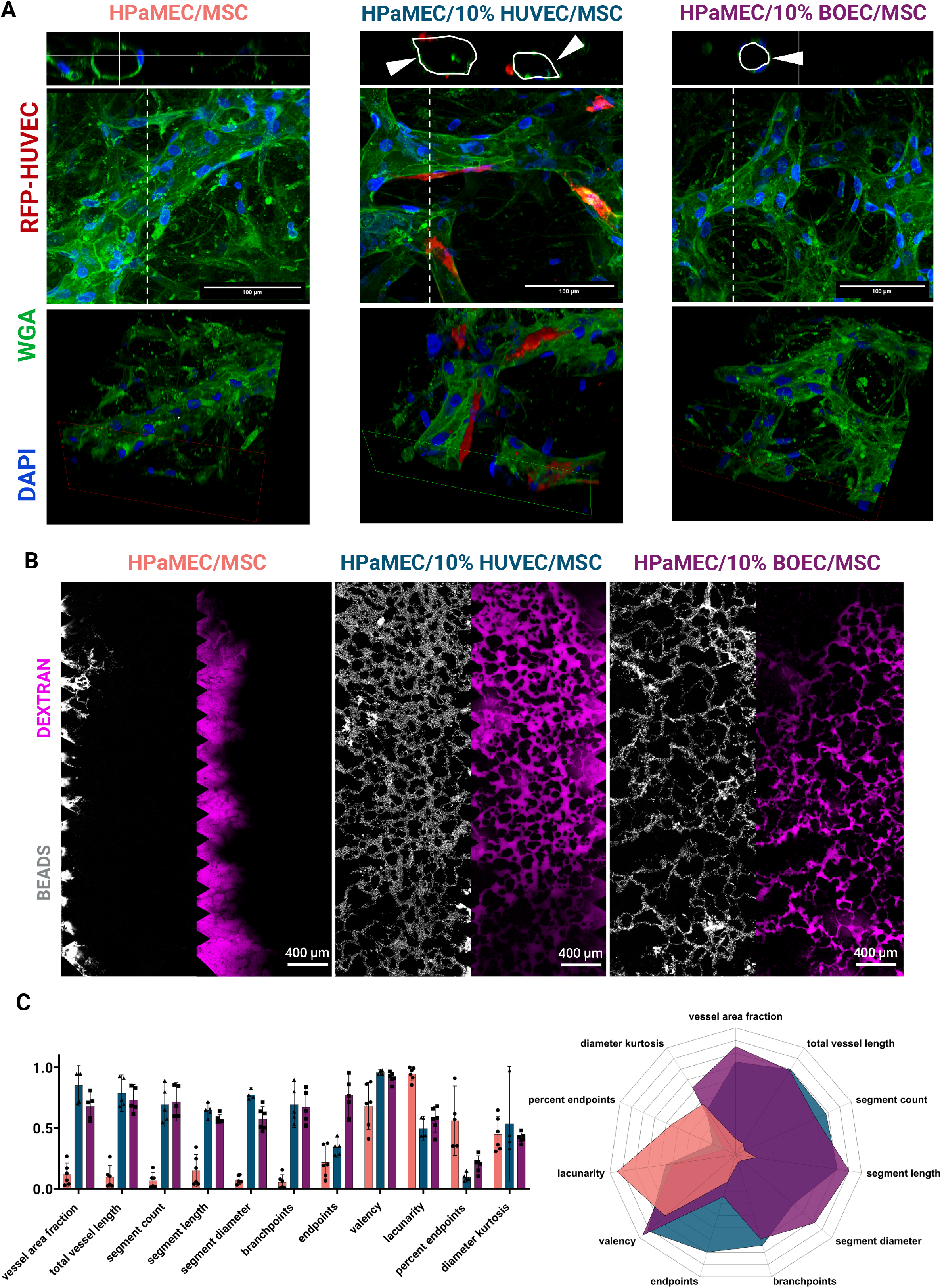
Potent endothelial sub-populations rescue network self-assembly. (A) Representative confocal images of HPaMEC and MSC, HPaMEC with 10% RFP positive HUVEC (red) and MSC; and HPaMEC with 10% BOEC and MSC co-culturing conditions. Top panel shows cross section of perfusable vessels for each condition with white arrowheads highlighting representative lumenised vessel structures. Middle panels show maximum-intensity projection (MIP) of immunofluorescently labelled constructs stained for cell membrane with WGA (green), and nuclei with DAPI (blue). Dashed lines indicate orthogonal views. Scale Bars 100 µm with 63x objectives. Bottom panels show three-dimensional reconstructions of above MIP. (B) Maximum intensity projection of 70kDa dextran (magenta) and 1 µm microbeads (white) perfusion through the vascular networks on day 5 for co-culturing of HPaMEC/MSC (n=6), co-cultured with 10% HUVECs (red) (n=5) and BOECs (n=6). Perfusion is performed unidirectionally from one medium channel to the other over the period of 5 minutes at 10x magnification. Scale Bar, 400 µm. (C) Quantitative network morphology analysis studying vessel area fraction, vessel length, segment count, segment length, segment diameter, branchpoints, endpoints, valency, lacunarity, percent endpoints, and diameter kurtosis between different endothelial conditions. All data are presented as means with 95% confidence intervals (CI).

### Immature EC sub-populations carry otherwise organotypic MVNs past 72-hour transcriptomic ‘stalling’

The addition of a small population of highly proliferative (HUVEC) or immature (BOEC) ECs rescued the self-assembly capabilities of a terminally differentiated HPaMEC cell line. To understand the mechanisms behind this, we sequenced bulk RNA from MVNs grown from MSCs and HPaMECs, MSCs and HPaMECs with 10% HUVECs, and MSCs and HPaMECs with 10% BOECs at days 0, 3 and 5. Notably, in contrast to earlier findings where MVNs were most similar according to their SC component (Fig. 2b), the EC-supplemented replicates clustered primarily by time point, with differences between the cell phenotype from day 0 and days 3 and 5 being most starkly segregated (Fig. 4a,b). Despite distinct EC provenances, all three MVNs converged in PCA by day 5. GO analysis revealed that the gene expression pattern of the three groups of networks evolved over time. The divergence in each type of engineered tissue revealed distinct pathways, which we conclude to have been fundamentally driven by the small populations of added BOECs or HUVECs (Fig. 4b and Supplementary Figure 3e).

**Figure 4.**
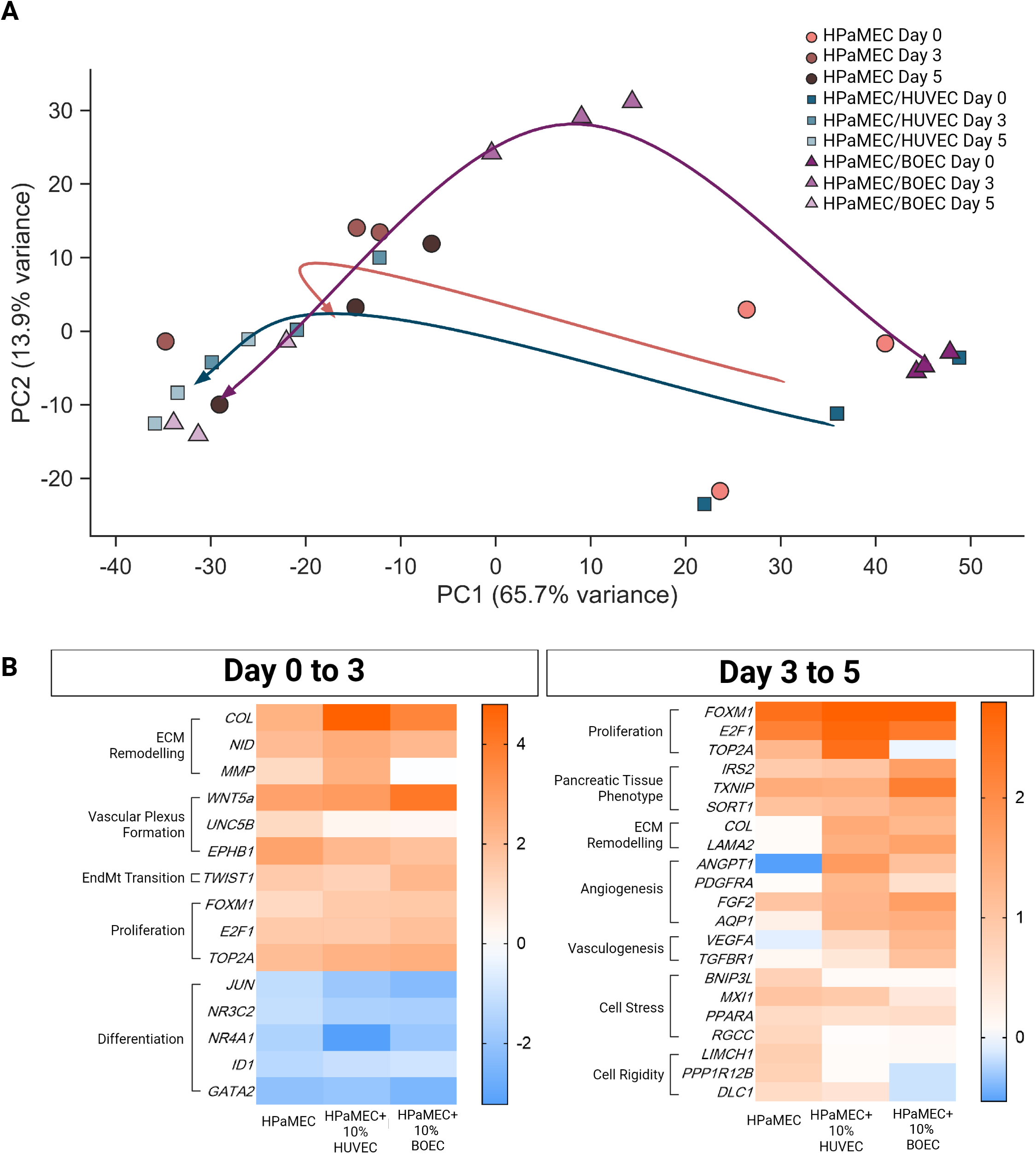
Potent endothelial cell sub-populations prevent terminal transcriptomic stalling. (A) Principal component analysis (PCA) of transcriptomic profiles of the organotypic endothelial conditions showing temporal progression and condition specific clustering. Each point represents two samples, with arrows inferring longitudinal developmental trajectories. (B) Heatmap of differentially expressed genes between day 0-3 and day 3-5 within each tissue condition, scale bars represent log fold change.

As before, all networks initially upregulated genes related to ECM, plexus formation, and EndMT. However, in comparison to the previous experiments that changed SC type, these chips that examined the addition of HUVECs to a set mix of HPaMECs and MSCs revealed a GO analysis more dominated by upregulated cell division, proliferation, and activity (FOXM1, E2F1, TOP2A), and downregulated pathways related to environmental cues and differentiation signalling (JUN, NR3C2, NR4A1, ID1, GATA2). By contrast, day 3 to 5 phenotype diverged significantly between HPaMEC/MSC only MVNs and those with added “rescue HUVECs”. Cells from all three groups showed a downregulation of genes related to growth and activity, while simultaneously upregulating genes associated with a pancreatic phenotype (IRS2, TXNIP, SORT1), suggesting network specialisation and consolidation. Interestingly, cells from HPaMEC/HUVEC and HPaMEC/BOEC networks displayed an upregulation of genes involved in angiogenic/tubulogenic pathways (ANGPT1, PDGFRA, FGF2, AQP1) and genes relating to wound healing and vasculogenesis (VEGFA, TGFBR1) respectively. Further, cells from HPaMEC/HUVEC networks also upregulated genes associated with native pancreatic ECM components, suggesting the emergence of a tissue-specific identity. The gene signatures of HPaMEC-only cultures, on the other hand, signalled cellular stress (BNIP3L, MXI1, PPARA, RGCC) and excess rigidity (LIMCH1, PPP1R12B, DLC1).

### Single-cell RNA-Seq highlights segregation in monolayer and on-chip cell phenotypes

Single cell RNA sequencing (scRNA-Seq) allows for the extraction of cellular subpopulations based on transcriptomic identity. We performed scRNA-Seq on cells extracted from MVNs grown from HPaMECs and MSCs and HPaMECs and MSCs with 10% RFP-HUVECs at day 0 and day 3 resulting in four libraries for analysis (Fig. 5a). Cellular identity was characterised by canonical marker expression (Fig. 5c-f). Quality control of sample libraries revealed tight clustering of samples by time point and cell type (Fig. 5b), and endothelial and stromal sub-populations were identified.

**Figure 5.**
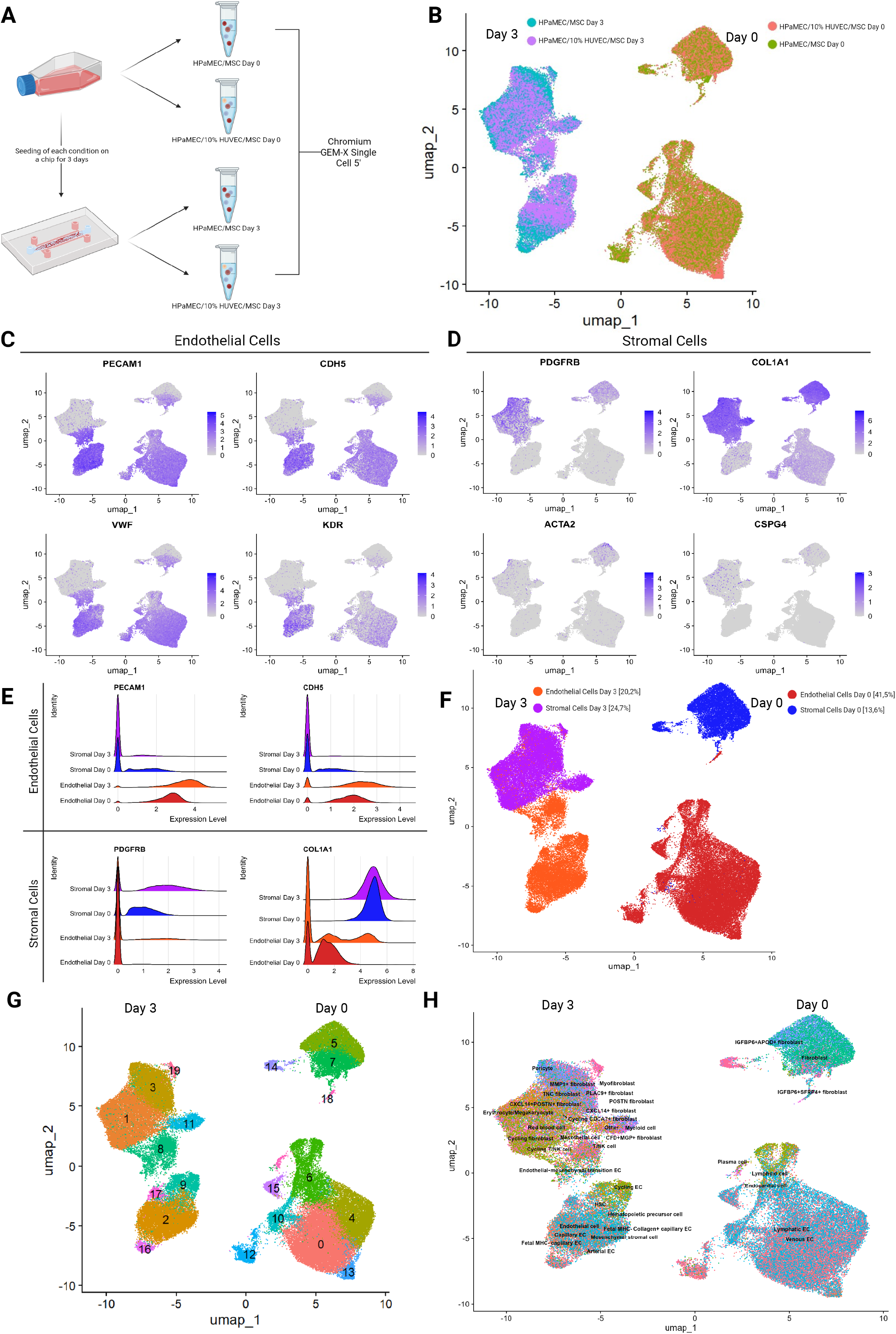
Single cell RNA sequencing reveals a distinct emergent phenotype in self-assembled networks. (A) Schematic overview of sample processing and single cell RNA sequencing extraction workflow. (B) UMAP visualisation of all cells from HPaMEC/MSC and HPaMEC/10% HUVEC/MSC flasks (day 0) and networks (day 3) after quality control filtering clustered by library (sample). (C, D) UMAP feature plots and (E) ridge plots of scaled RNA expression of canonical marker genes for endothelial and stromal cells, and (F) UMAP visualisation and quantification and clustering of all cells based on said markers. (G) UMAP visualisation of 20 cell sub-clusters across four libraries. (H) UMAP visualisation and clustering of all cells using Pan-Human Azimuth cell characterisation. Data represents n=2 technical replicates per condition. Cell clustering was performed using Seurat and cell characterisation using AzimuthAPI.

Endothelial-associated clusters were defined by high expression of CDH5 (VE-cadherin), PECAM1, KDR (VEGFR2), and VWF, while stromal-associated clusters were identified by COL1A1, PDGFRB, CSPG4 and ACTA2 (Fig. 5c,d). Pan-Human Azimuth annotation of sample libraries identified more diverse cell populations in day 3 networks when compared to day 0 cells, as well as the emergence of cell sub-populations such as capillary and arterial ECs, EndMT ECs and MSCs all positive for canonical EC markers (Fig. 5h).

### Self-assembling vessel networks feature highly specialised EC sub-populations

Further analysis of a single sample library representing the day 3 phenotype of a successfully self-assembled vascular network (HPaMEC/10% HUVEC/MSC) resolved five transcriptionally distinct endothelial sub-populations corresponding to discrete stages of vascular morphogenesis (Fig. 6a, b). Tip cells (EC1) were enriched for programmes such as VEGF-responsive migration and chemotaxis, stalk cells (EC3) were undergoing active mitosis, and phalanx cells (EC5) upregulated negative locomotion and barrier function. Additionally to these groups well-established to support network formation was another set of endothelial sub-populations adopting what was previously thought to be stromal cell behaviours. EC2 was a population of cells undergoing EndMT, a well described process, but in considerable numbers (Fig. 6c,d, Supplementary Figure 4). Additionally, CellChat revealed complex and multidimensional co-regulatory signalling pathways between EndMT and tip cell sub-populations (Fig. 6e, Supplementary Figure 5). Tip cells showed an upregulation of genes encoding for WNT and ANGPT ligands while EndMT cells showed a corresponding upregulation of non-canonical WNT and ANGPTL signalling pathways. Tip cells were also found to be enriched for basement membrane (BM) components that are typically found on the abluminal surface such as laminin and perlecan, whereas EndMT cells secreted adluminal ECM elements, such as Collagen, Fibronectin, Tenascin, and Periostin. Critically, EC4 was identified as an EC sub-population enriched for cell-cell and cell-substrate adhesion, suggesting the emergence of a quiescent pericyte-like niche likely differentiated from the HUVEC population which bears progenitor-like features.

**Figure 6.**
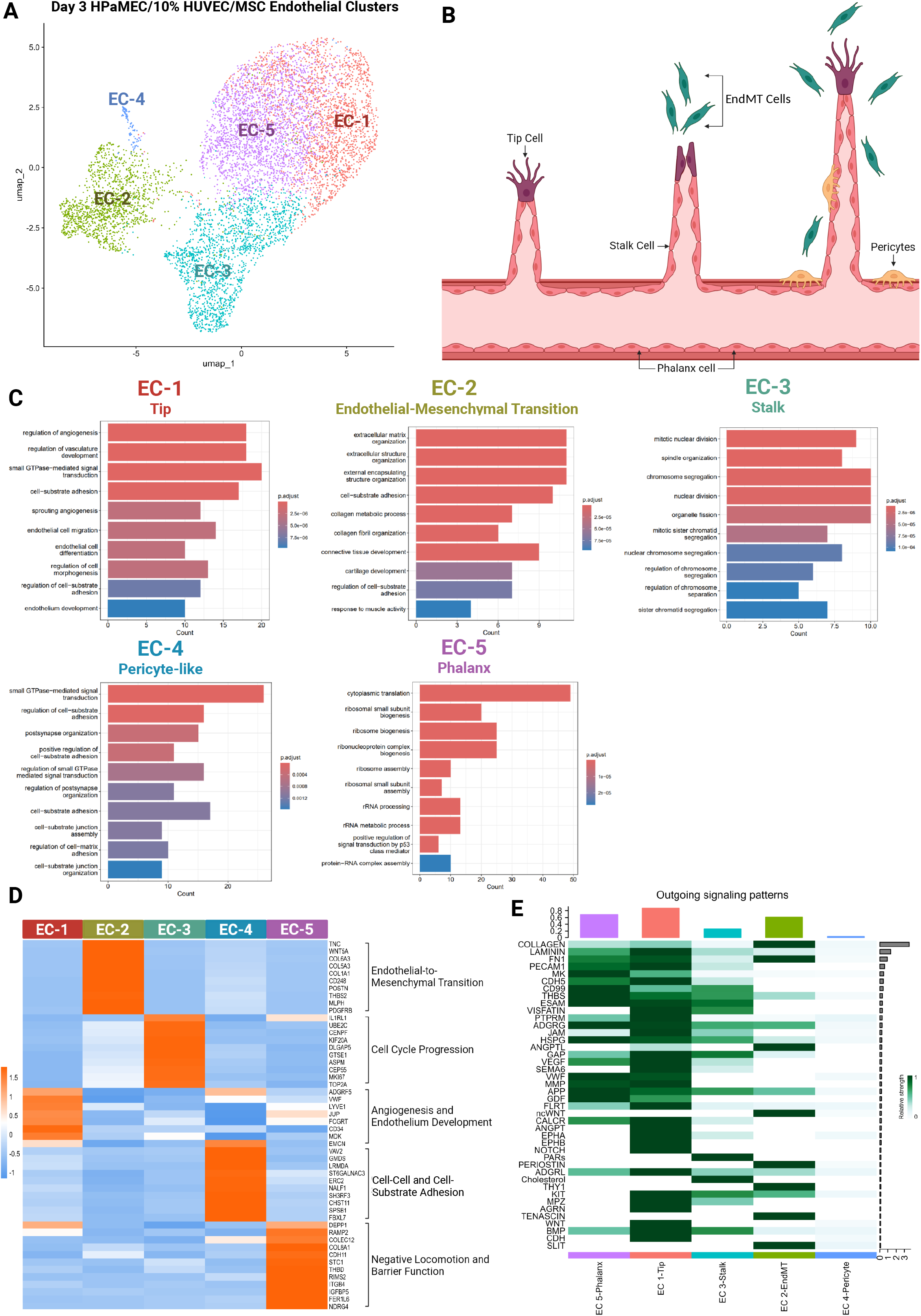
Successfully self-assembled networks feature five specialised endothelial cell sub-populations. (A) UMAP visualisation of the endothelial cell portion of HPaMEC/10% HUVEC/MSC networks on day 3 identified by canonical markers and clustered by gene expression. (B) Schematic overview of vasculogenesis in a developing self-assembled network. (C) Functional GO enrichment analysis of upregulated differentially expressed gene sets in each EC sub-group. Bar plots show representative biological processes, bar size reflects gene count and colour indicates statistical significance (adjusted p-value). (D) Heatmap of log2 fold change of marker genes expressed by endothelial cell sub-clusters. (E) Heatmap of all outgoing signalling among endothelial cell sub-clusters. Seurat’s FindAllMarkers was used to identify markers and CellChat was used to identify signalling.

### Rescued network HUVECs potentiate angiogenesis via the FLRT2-UNC5B axis

We next investigated what drives failure in self-assembling networks of terminally differentiated ECs, and via what mechanisms progenitor-like ECs are able to rescue this process. On their own, HPaMEC/MSC networks demonstrate a markedly different signalling footprint compared to HPaMEC/10% HUVEC/MSC, even within EC subpopulations alone. Notably absent were both phalanx and pericyte-like subpopulations, implying a complete lack of a stabilised network niche (Fig. 7b). Furthermore, tip cell identity was disordered and less homogeneous, with contradictory gene signatures emerging within the same cell population (angiogenesis and barrier function, distal and proximal elements of BM (Fig. 7c, Supplementary Figure 7). In brief, network developmental failure is associated with a breakdown in coherent intercellular signalling. We used Souporcell to separate scRNA-Seq datasets by cell type via the donor genome, allowing us to remove MSC contaminant from EC groups and isolate the critical subpopulation of RFP-positive HUVECs from HPaMECs (Fig. 7a). Despite being represented disparately across each EC sub-group, HUVECs were over-represented within the EndMT population. Additionally, though most HUVEC subgroups were not meaningfully distinct from the co-loaded HPaMECs, pericyte-like HUVECs specifically expressed one unique signalling axis: FLRT2-UNC5B (Fig. 7f,g and Supplementary Figure 8). This pathway could not be detected at meaningful levels in HPaMEC/MSC networks. This pathway is well known to regulate embryonic endothelial development and angiogenesis, and notably, knockdown of either FLRT2 or UNC5B leads to the destruction of vessel formation in zebrafish^55,56^ in a manner that is strikingly visually comparable in an animal model to these NAM networks.

**Figure 7.**
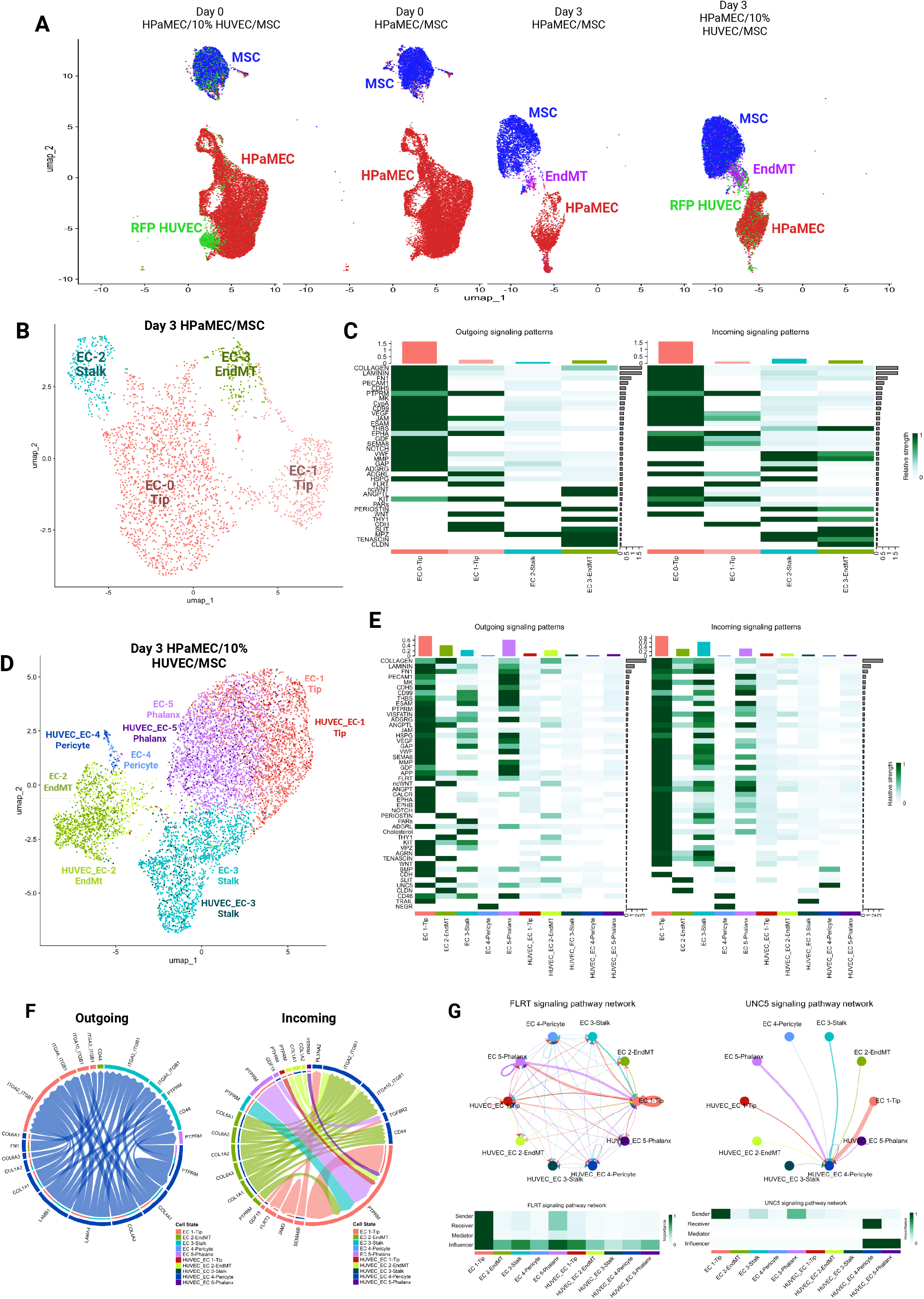
FLRT2-UNC5B potentiates angiogenesis in rescued networks. (A) UMAP visualisation of each sample library separated by time point and clustered by Souporcell-derived donor provenance. Provenance sorting described in methods. (B) UMAP visualisation of the endothelial cell portion of HPaMEC/MSC networks on day 3 identified by canonical markers and clustered by gene expression. (C) Heatmap of all outgoing and incoming signalling among endothelial cell sub-clusters in HPaMEC/MSC networks. (D) UMAP visualisation of the endothelial cell portion of HPaMEC/10%HUVEC/MSC networks on day 3 identified by canonical markers and clustered by gene expression and donor provenance. (E) Heatmap of all outgoing and incoming signalling in the aforementioned. (F) Chord plots of top predicted ligand-receptor interactions in the HUVEC-Pericyte subpopulation in HPaMEC/10%HUVEC/MSC networks on day 3. (G) Cell-cell communications of FLRT and UNC5 signalling in HPaMEC/10%HUVEC/MSC networks on day 3. Souporcell was used to identify donor provenance and CellChat to identify signalling between cell sub-populations.

## Discussion

Vascularised organ-on-chip NAM platforms require not only robust self-assembly and perfusability, but also faithful recapitulation of structure, organotypic function and transcriptional identity. For a NAM to be adopted it must be substantiated for its given context of use (CoU). In the past decade there have been great advances in the engineering aspects of perfusable OoC MVNs^57^, but arguably less implementation of these platforms at scale for physiological or drug testing. It is therefore important to establish further validatory data for their broader application. Here we demonstrate that both stromal and endothelial identity play a deciding role in MVN development and architecture. These data establish that cell sources often treated as interchangeable in OoC models are, in fact, quantifiably distinct and should be accounted for in the CoU. We also show that the self-assembly capability of terminally differentiated organotypic ECs may be rescued, and that engineered microvasculature can accurately model *in vivo* vascular function to the receptor-ligand level.

SCs in MVN experiments are often selected pragmatically rather than biologically^6^, however, we find that SCs longitudinally impart key characteristics to self-assembled MVNs. HUVEC/MSC networks formed dense vascular beds with variably sized thick vessels, and active proliferative and mural-associated pathways, consistent with the relatively immature phenotype of both cell types. HUVEC/nHLF networks formed thinner vessels, and the fibroblasts significantly upregulated more pathways associated with integrin-mediated signalling – a feature that has been reported in mono-culture^46^. The disintegration of HUVEC/HPF networks was corroborated by a transcriptomic phenotype dominated by sustained metabolic activation, inflammatory signalling, and downregulation of NO pathways. Rather than converging towards a stabilised microvascular phenotype, these networks were morphologically and transcriptionally dysregulated. Given the specialised and heterogeneous nature of pancreatic stromal niches *in vivo* ^58^, it is plausible that isolated pancreatic fibroblasts, removed from their native microenvironment, struggle to provide balanced angiocrine cues. Importantly, these findings highlight that organ-matched SCs do not automatically confer organotypic behaviour onto engineered networks, and that the frequent interchangeability of MSCs, nHLFs, and other fibroblasts in OoC literature introduces under-recognised variability across platforms.

It has long been known that ECs of a given tissue specialise according to their niche during angio- and vasculogenesis. However, due to the experimental impenetrability of animal models, many of these nuances have barely been characterised. Taking advantage of new approach methodologies for growing vascular tissues *in vitro*, and the pipeline for single-cell sequencing, it is now possible to interrogate some of these developmental events in a new way. We are able to identify in our networks the presence of tip, stalk and phalanx EC sub-populations canonical to the process of vascularisation, but also the presence of EndMT-ECs, and the unexpected transition of some ECs into pericyte-like cells. Each of these EC sub-populations is involved in constant communication and co-regulation with one another, and indeed this dialogue appears essential for proper engineered vascular development, as when it is incoherent (as with HPaMEC networks), self-assembly fails.

The exceptional vasculogenic capacity of HUVECs, while experimentally advantageous, may mean they are less relevant for particular CoUs. However, our work with HPaMECs suggests that terminally differentiated primary ECs are very limited in their OoC vasculogenic potential ^59^. We found that this may be rescued with the addition of a small subpopulation of immature ECs (either HUVECs or BOECs) to fully restore network formation in HPaMEC-based co-cultures. It may seem intuitive that the reason “adult” primary ECs are less angiogenic than HUVECs is because they fail to sprout or initiate angiogenesis, but this model suggests something different; what HPaMEC networks lack are the proper mechanisms to tune angiogenesis. HPaMECs are unable to suppress sprouting, which hinders lumen formation and stabilisation, resulting in “frayed” and “leaky” vessels on imaging. Our specific findings, that networks with added HUVECs potentiate angiogenesis via the ligand-receptor pair FLRT2-UNC5B, also validate the chip as a promising NAM. Zebrafish and murine models have already confirmed that FLRT2 and UNC5B are regulators of angiogenic development. This human NAM model takes those findings further by identifying a specific subpopulation of ECs that take on a pericyte-like phenotype and express UNC5B to respond to FLRT2. Future work will be to characterise this pathway for regenerative applications, as well as developing a pipeline for even better organotypic vasculatures for defined CoUs.

Taken together, these data support three principal conclusions regarding the application of microvascular NAMs; that detailed cell provenance is essential for defining the CoU, that the self-assembly capability of terminally differentiated organotypic endothelial cells can be rescued *in vitro*, and that chip-grown vasculature is able to recapitulate and extend findings from animal models.

## Methods

### Cell Culture

Human Umbilical Vein Endothelial Cells (HUVECs) expressing red fluorescent protein (RFP) (Angioproteomie), human Blood Outgrowth Endothelial Cells (BOECs) provided by the Ahmetaj-Shala Group (isolated from blood donated from human healthy donors as previously described^27^) and primary Human Pancreatic Microvascular Endothelial Cells (HPaMECs) (2BScientific) were maintained in EGM-2 (Endothelial Cell Growth Medium-2 BulletKit, Lonza), prepared using the supplements as directed by the manufacturer, with the exception of Foetal Bovine Serum (FBS) (ThermoFisher), which was added at a higher concentration (10% v/v). Mesenchymal Stem Cells (MSCs) (PromoCell), Normal Human Lung Fibroblasts (nHLF) (Lonza) and Human Pancreatic Fibroblasts (HPF) (2BScientific) were cultured in FGM-2 Fibroblast Growth Medium-2 BulletKit (Lonza). All cell lines were used between passages 3-9 and cultured in a humidified incubator at 37 °C and 5% CO_2_ with media replacement every second day. Upon reaching near-confluence, cells were dissociated using TrypLE (ThermoFisher).

### Microfluidic Device Fabrication

Computer-aided design (CAD) files of the chip designs were created using Inventor 2024 (Autodesk) and inspired by the 3-channel design divided by posts as described in literature ^36,60^. The printing of the photomask was outsourced to Micro Lithography Service and used to manufacture SU-8 micropatterned silicon wafer masters. Silicon wafers (MSE Supplies) were cleaned with 99% isopropanol and oxygen plasma treated (Harrick) at 0.5 Torr O_2_ for 5 minutes before spin coating with a negative photoresist SU-8 2100 (Kayaku). Wafers were spun at 500 rpm for 20 seconds followed by a second spin at 1500rpm for 40s for even dispersion of the resist up to a height of 200µm. This was followed by a soft bake at 65°C for 6 minutes and another at 95 °C for 37 minutes. SU-8 coated wafers with the desired photomask were exposed to UV photolithography with a UV-KUB 3 mask aligner (KLOE). The crosslinked wafer was post-baked at 65°C for 5 minutes, followed by 95 °C for 13 minutes. The wafers were soaked in fresh SU-8 developer for 5 minutes three times. Finally, the masters were rinsed using isopropanol to remove any excess SU-8 photoresist.

Polydimethylsiloxane (PDMS Sylgard 184) was mixed at a ratio of 10:1 for the device top layer. The mixture was degassed for 30 minutes, poured into the mould and cured overnight in an oven at 65°C. The microfluidic chips were cut individually, and media ports were punched using a 1.5mm biopsy punch. PDMS chip devices were bonded to glass slides (VWR) through plasma treatment using O_2_ at 0.5 Torr for 5 minutes (Expanded plasma cleaner, Harrick Plasma) and cured at 90°C on a hot plate for 15 minutes.

### Device Seeding

Different types of ECs and SCs were maintained until reaching confluency and dissociated using TrypLE (ThermoFisher). Co-cultures were resuspended at 10x10^6^ cells/mL of ECs and 1.5x10^6^ cells/mL of SCs in EGM-2 medium and 4U/mL concentration of thrombin (Sigma-Aldrich). For HPaMEC chips with added populations of HUVECs or BOECs, 10 percent of the latter cell type was added to 90 percent of the former and the population of MSCs was unchanged. For each chip device, a volume of 10 µL of cell suspension with thrombin was combined with 10 µL of fibrinogen (Scientific Laboratory Supplies) at 6 mg/mL and injected through the loading port of the central channel. The microfluidic chips were incubated at 37°C for 30 minutes prior to adding EGM-2 medium in the media channels to allow for fibrin gelation. Chips were allowed 24 hours initially under equivalent pressure (200:200µL media) and subsequently medium was changed daily under an alternating pressure gradient of 600:200 µL.

### Microvascular Network Perfusion

To assess the functional perfusability of the formed vascular networks, medium was removed from the external feeding channels at day 5 and fluorescent polystyrene beads of 1 µm in diameter suspended in DPBS and FITC dextran 70kDa (ThermoFisher) solution was injected to assess the perfusability of the grown networks in the microfluidic devices. The chip was imaged using an SP8 Leica confocal laser scanning microscope with a 10x objective as well as a Leica Stellaris 8 with a 63x objective. Z-stacks were taken at 5 µm step size and about 10-15 xy-slices acquired per image.

### Image Analysis

Basic image processing was performed using Fiji (ImageJ). Network morphology was analysed using a bespoke MATLAB algorithm available upon request. Briefly, maximum intensity projections of z-stack tilescans acquired as described above were binarised with manual thresholding and processed to remove artefacts and fill holes. Resultant images were skeletonised, branchpoints and endpoints were identified, and images were segmented to create a mask of the vessel network (Supplementary Figure 1a). Finally, all statistics generated via this process were exported to Excel for downstream analysis.

### Immunostaining

For staining, chips were fixed for an hour with 4% methanol-free PFA in DPBS (ThermoFisher) at room temperature by hydrostatic pressure-driven perfusion. Following fixation, cells were washed with PBS for 30 minutes. For cell surface staining, a WGA (ThermoFisher) conjugate (working solution of 5 µg/mL in DPBS) was applied, using the same method, and incubated for 20 minutes at room temperature, protected from light. Excess WGA was removed by washing once with DPBS. Nuclei were stained with DAPI (working solution of 1 µg/mL in DPBS) for 30 minutes, followed by DPBS washes. Channels were imaged directly using a confocal microscope. This protocol enabled simultaneous visualisation of cell membranes (WGA) and nuclei (DAPI) within the microfluidic system.

### Bulk RNA Sequencing

We performed bulk RNA sequencing to investigate the gene expression phenotypes of the different seeding populations. The microfluidic chip design was adjusted and upscaled to allow extraction of engineered tissue from the central channel. The design of the chip was made using CAD software Inventor 2024 (Autodesk) and 3D printed with VeroClear in an Objet500 Connex3 (Stratasys). The design changes consisted of three square-profile channels with a height of 400 µm, separated by down-step extrusion (Fig. 2a). To retrieve engineered tissue from the microfluidic chips, the glass slide was carefully detached from the PDMS device using a scalpel. The cell-laden fibrin was collected and suspended in an RNA protection reagent solution. RNA was extracted using Monarch Total RNA Miniprep Kit (New England Biolabs), following manufacturer’s instructions, with the exception that the Proteinase K incubation time was extended to 6 minutes. The RNA integrity number (RIN) was checked with 2100 Bioanalyser (Agilent Technologies Co), the RNA samples with RIN > 8 were sent to Novogene (Cambridge) for subsequent sequencing. Libraries of the samples were sequenced using an Illumina Novaseq Xplus platform with PE150 (Novogene), following standard Illumina protocols by Novogene.

### Pre-processing of bulk transcriptome data

ENSEMBL gene IDs were converted into gene symbols using the mapIds() function of the R/Bioconductor package AnnotationDbi (v1.60.0). Only genes with a gene symbol and with a total sum of reads across all conditions greater than or equal to 20 were kept for further analysis. Gene expression values were normalised with the R/Bioconductor package DESeq2 (v1.38.3) and only false discovery rate-adjusted P values with a value of less than 0.05 were considered statistically significant. Log base 2 fold change-transformed FPKM values were used for heat maps, and FPKM values were used for PCA. The labelling of differentially expressed genes (p-value < 0.05) across the different groups of experimental conditions was acquired with the non-parametric Kruskal-Wallis rank test from normalised gene expression values using the col_kruskalwallis function of the CRAN package matrixTests (v.0.1.9.1). The ontological enrichments were performed from the lists of differentially expressed genes with the R/Bioconductor package clusterProfiler (v.4.6.0) (*p*-value < 0.05; Benjamini-Hochberg *q*-value < 0.05).

### Sample Collection from microfluidic chip for single-cell RNA sequencing

Based on the work from Carrion et al.’s group ^61^, a protocol was optimised for our microfluidic devices. Nattokinase powder was dissolved in DPBS at a concentration of 100 FU/mL and incubated at 37°C. The solution was filter sterilised using a 0.2 µm pore-sized filter before treatment in the chip. Microfluidic chips were opened using the previously described method and washed with DPBS before the samples were collected. The engineered tissue was then incubated with 1:1 ratios of nattokinase solution for 15 minutes at 37°C. Cell solution was homogenised and centrifuged at 300 x g for 5 minutes. Cell pellet was resuspended in EGM-2 media and filtered through a 37 µm pore-sized filter. Cell viability was checked and samples above 85% were stored at 4°C for immediate processing for scRNA-seq analysis.

### Single-Cell library preparation and sequencing

ScRNA-Seq libraries were prepared from isolated cells by the NIHR Imperial BRC Genomics Facility using a 10X Genomics Chromium X (10X Genomics) according to manufacturer’s instructions. Libraries were sequenced using Illumina NextSeq 2000 sequencers.

### Basic processing and quality control of SC RNA Seq data

Raw demultiplexed sequencing data were mapped to the human genome (GRCh38), quantified using cellranger-arc (10X Genomics) and loaded into an R environment (R v4.3.3), using the Seurat single cell analysis package v5.1.0. To retain only high quality datasets and cells, low quality cells and potential cell clumps were removed, i.e. cells with low or extremely high transcript counts (<500 or > 10,000 UMIs/cell), or high counts of mitochondrial genes (>10%). The resultant database contained 28638 genes across 67797 cells with the median UMI count being 5634 and the median number of unique genes per cell being 2558. For all libraries when performing UMAPs, the clustree package was used to visualise group lineage across varying resolutions and select the ideal resolution. Generally, robust clusters were identified at the resolution 0.4. Endothelial cells were identified using the markers PECAM1, CDH5, VWF, and KDR, and stromal cells primarily using the markers PDGFRB and COL1A1, though ACTA2 (smooth muscle cells) and CSPG4 (pericytes) were also included for interest. For isolated analysis, endothelial cell portions in all network types were segregated based upon a combination of their canonical marker expression and generic clustering. GO was run on isolated cell portions in the same manner aforementioned, and heat maps were generated using the FindAllMarkers function. CellChat v2.2.0.9001 was used to identify and analyse ligand receptor pairs in all isolated datasets with a minimum cell number of 10. Finally, Souporcell was used to identify and isolate cell provenance based on donor genotype, and all HPaMEC datasets were run with a total donor number of 2 and HPaMEC/10%HUVEC with a total donor number of 3. Cells expressing both endothelial and stromal canonical marker genes in both cases that could not be sorted into a single donor genotype were characterized as EndMT.

### Statistical Analysis of Imaging Data

All data are presented as means with 95% confidence intervals (CI). Differences between conditions were analysed by one-way ANOVA. Where ANOVAs showed significant effect, post-hoc pairwise comparisons between all three conditions were performed within each parameter using Tukey’s multiple comparisons test. Parametric assumptions were assessed on the ANOVA residuals, normality was evaluated using the Shapiro-Wilk test. Equal variances between groups were evaluated using Brown-Forsythe and Bartlett’s test and visual inspection of QQ plots. All statistical analyses were performed using GraphPad Prism (v11.1.0(224), GraphPad Software), and MATLAB (R2026b; MathWorks), as appropriate. Results were calculated by averaging the mean values of at least n=5 microfluidic devices. Statistical significance was defined as p < 0.05. For all statistics, p-value numbers were graphically depicted as n.s. = p > 0.05, * = p < 0.05, ** = p < 0.01, *** = p < 0.001 and **** = p < 0.0001 in the results.

## Ethics Statement

Ethical approval was obtained from (i) Imperial College Research Ethics Committee (19IC5372) and (ii) REC Wales approval (17/WA/0161) as per the Declaration of Helsinki. Informed written consent was given prior to the inclusion of subjects in the study. All cells were used between passages 3 and 9. A single BOEC donor was used in this study.

## Acknowledgements

This research was supported by the Type 1 Diabetes Grand Challenge, a Diabetes UK RD Lawrence Fellowship to VS, and NC3Rs, and in part by a Royal Society Wolfson Research Merit Award to PP. Imaging was supported by the Imperial College London and Leica Microsystems Imaging Hub. The Imperial BRC Genomics Facility provided resources and support that contributed to the research results reported in this paper. The facility is supported by NIHR funding to the Imperial Biomedical Research Centre. We thank members of the Au, Pantazis, van Batenburg-Sherwood and Ahmetaj-Shala groups for their experimental support and valuable discussions. We also extend our thanks to the blood donors whose selfless dedication to medical research made this work possible. Schematic diagrams were created with BioRender.com.

## Author Contribution

L.A.O., B.H and V.S. designed the research; L.A.O., B.H. performed the experiments; L.A.O. developed and optimised methodology; L.A.O. and B.H. analysed and interpreted the data; B.H. performed bioinformatic analysis; V.S. acquired the funding; B.A.S. provided patient samples; L.A.O., B.H. wrote and edited the article with input from V.S.; K.S., Y.Y., M.N., X.X., and M.W. provided experimental support; P.P., S.A., J.V.B.S., B.A.S. and I.A. provided scientific insight and resources. All authors reviewed, discussed and edited the manuscript.

## Declaration of Interests

The authors declare no competing interests.

---

**Supplementary Figure 1. Workflow for quantification of microvessel networks using a bespoke algorithm.** (A) Detailed workflow of microvessel network quantification. Microfluidic chips were imaged on Day 5, once the network was fully perfused. The devices were perfused with 70 kDa dextran and imaged using a confocal microscope, and the resulting images were processed using ImageJ. The processed images were then quantified using a laboratory-developed algorithm, which generated skeletonised images and enabled statistical comparison of different microvessel network conditions through the described image processing steps. (B) Quantitative network morphology analysis studying vessel area fraction, segment length, segment diameter, valency, lacunarity, percent endpoints, and diameter kurtosis between different stromal conditions. All data are presented as means with 95% confidence intervals (CI). *p<0.05; **p<0.01; ***p<0.001; ****p<0.0001

**Supplementary Figure 2. Longitudinal differential gene expression from different stromal cell types.** (A) Volcano plots showing differential gene expression in adipose-derived, lung and pancreatic stromal cell conditions. Genes meeting the defined thresholds (adjusted *P* < 0.05 and |log₂ fold change| ≥ 0.5) are highlighted in red, while genes not meeting both criteria are shown in grey. (B) Functional GO enrichment analysis of differentially expressed gene sets. Dot plots show representative longitudinal biological processes and pathways enriched in the different tissue conditions. Dot size reflects gene count, and colour indicates statistical significance (adjusted p-value). Data represents n=6 biological replicates per condition. Differential expression analysis was performed using DESeq2; enrichment analysis was performed using clusterProfiler. Statistical thresholds and normalisation steps are described in methods.

**Supplementary Figure 3. Organotypic cell microvessel network architecture can be modulated with the aid of other endothelial cell lines.** (A) Quantitative network morphology analysis studying vessel area fraction, vessel length, segment count, segment length, segment diameter, branchpoints, endpoints, valency, lacunarity, percent endpoints, and diameter kurtosis between different endothelial conditions. All data are presented as means with 95% confidence intervals (CI). *p<0.05; **p<0.01; ***p<0.001; ****p<0.0001 (B) Maximum intensity projection of 70kDa dextran (magenta) and 1 µm microbeads (white) perfusion through the vascular networks on day 5 for co-culturing of HPaMEC/MSC. Perfusion is performed unidirectionally from one medium channel to the other over the period of 5 minutes at 10x magnification. Scale Bar, 400 µm (left) and 100 µm (right). (C) HPaMEC with 10% RFP positive HUVEC (red) and MSC with dextran on day 3. HPaMEC with 10% BOEC and MSC co-culturing conditions at days 2 and 4.(D) Volcano plots showing differential gene expression in endothelial cell conditions. Genes meeting the defined thresholds (adjusted *P* < 0.05 and |log₂ fold change| ≥ 0.5) are highlighted in red, while genes not meeting both criteria are shown in grey. (E) Functional GO enrichment analysis of differentially expressed gene sets. Dot plots show representative longitudinal biological processes and pathways enriched in the different tissue conditions. Dot size reflects gene count, and colour indicates statistical significance (adjusted p-value). Data represents n=6 biological replicates per condition. Differential expression analysis was performed using DESeq2; enrichment analysis was performed using clusterProfiler. Statistical thresholds and normalisation steps are described in methods.

**Supplementary Figure 4. Successfully self-assembled networks feature five specialised endothelial cell sub-populations.** Functional GO enrichment analysis of downregulated differentially expressed gene sets in each EC sub-group. Bar plots show representative biological processes, bar size reflects gene count, and colour indicates statistical significance (adjusted p-value).

**Supplementary Figure 5. Cell-cell communications in tip and phalanx cells in the endothelial portion of HPaMEC/10%HUVEC/MSC networks on day 3.** Circle plots of cell-cell communications of VEGF, WNT, ANGPT, Laminin, HSPG, MMP, THBS, VWF and CD99 signalling.

**Supplementary Figure 6. Cell-cell communications in stalk, EndMT and pericyte-like cells in the endothelial portion of HPaMEC/10%HUVEC/MSC networks on day 3.** Circle plots of cell-cell communications of PARs, Cholesterol, BMP, ncWNT, ANGPTL, Periostin, Collagen and FN1 signalling.

**Supplementary Figure 7. Cell-cell communications in all cell subpopulations in the endothelial portion of HPaMEC/MSC networks on day 3.** Circle plots of cell-cell communications of PTPRM, WNT, PARs, NOTCH, ncWNT, ANGPTL, Tenascin, Periostin, VEGF, ADGRG, VWF, MMP, Collagen and Laminin signalling.

**Supplementary Figure 8. Donor provenance in the endothelial portion of HPaMEC/10% HUVEC/MSC networks on day 3 and Ligand-receptor interactions in RFP-pericyte-like cells.** (A) UMAP visualisation of cells from the endothelial portion of HPaMEC/10%HUVEC/MSC networks on day 3 clustered by cell type and separated by donor provenance. Souporcell was used to identify donor provenance. (B) Circle plots of predicted ligand-receptor interactions in the RFP-pericyte-like subpopulation of the endothelial portion of HPaMEC/10% HUVEC/MSC networks on day 3. (C) Circle plots of all predicted ligand-receptor interactions in the aforementioned.

## Notes

### Competing Interest Statement

The authors have declared no competing interest.

