## Supplementary Figures for "Engineered microvessel networks – Longitudinal transcriptomic profiling validates chip-grown tissues as a robust model for studying vasculature"

### Supplementary Figure 1

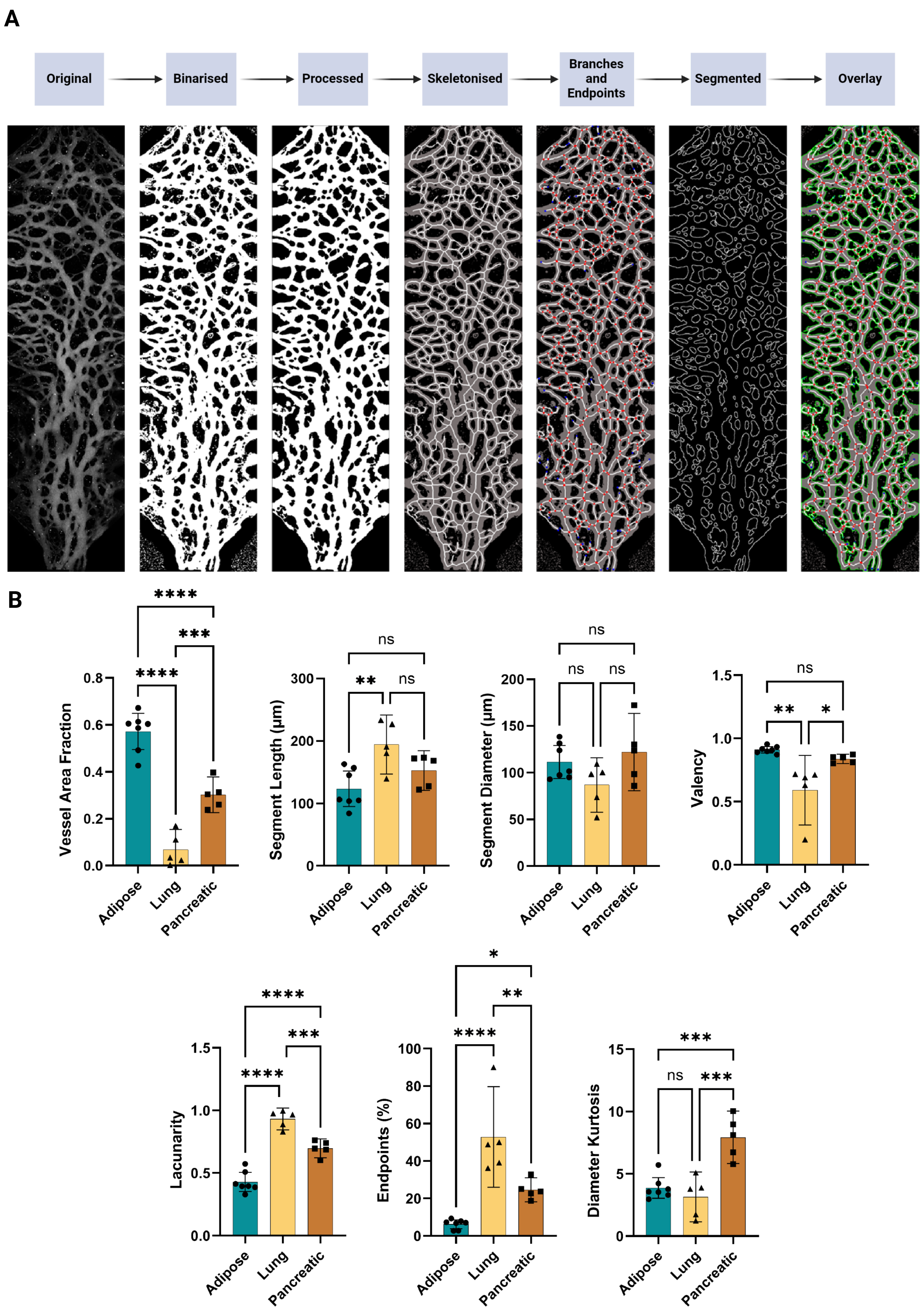

### Supplementary Figure 2

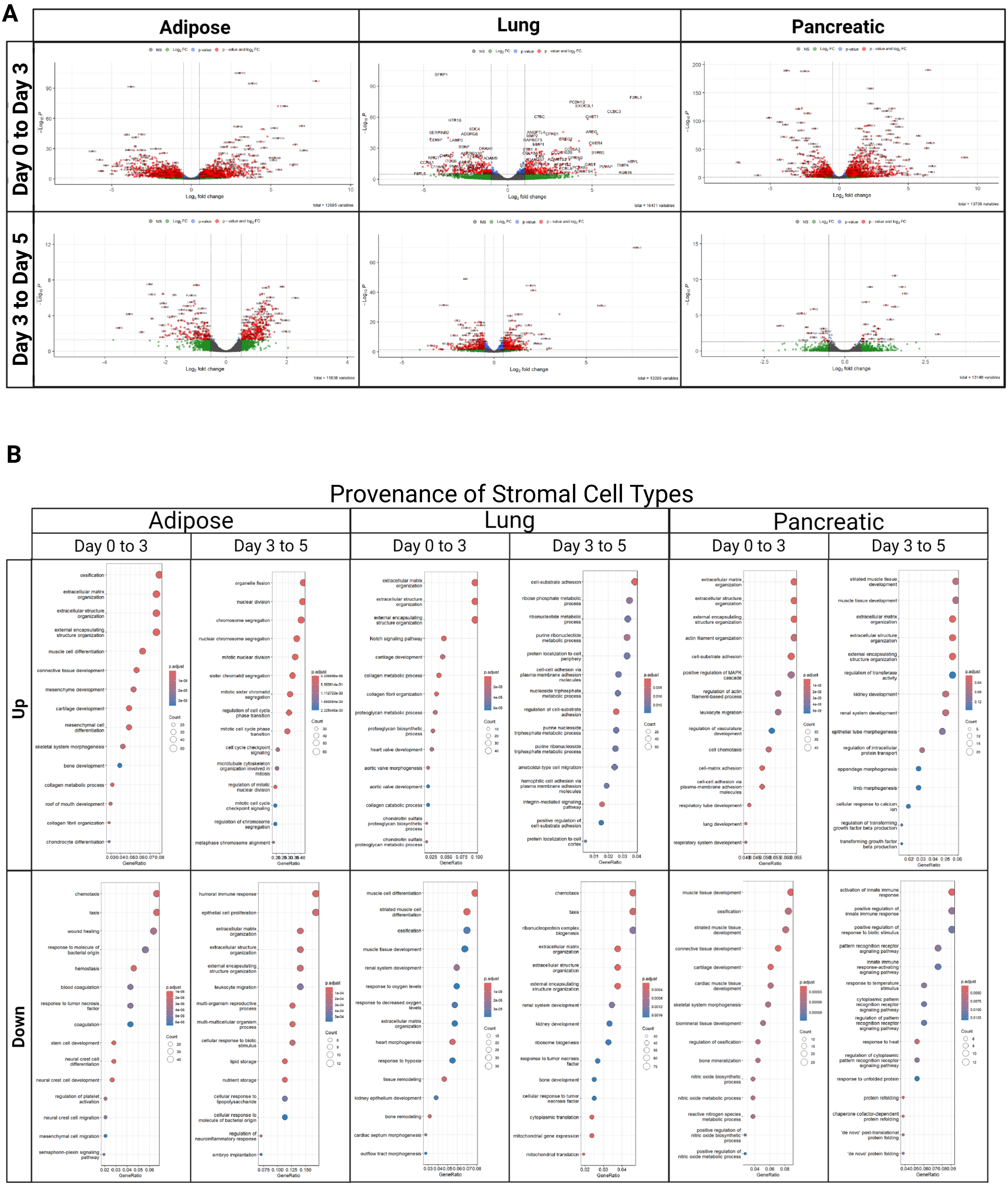

### Supplementary Figure 3

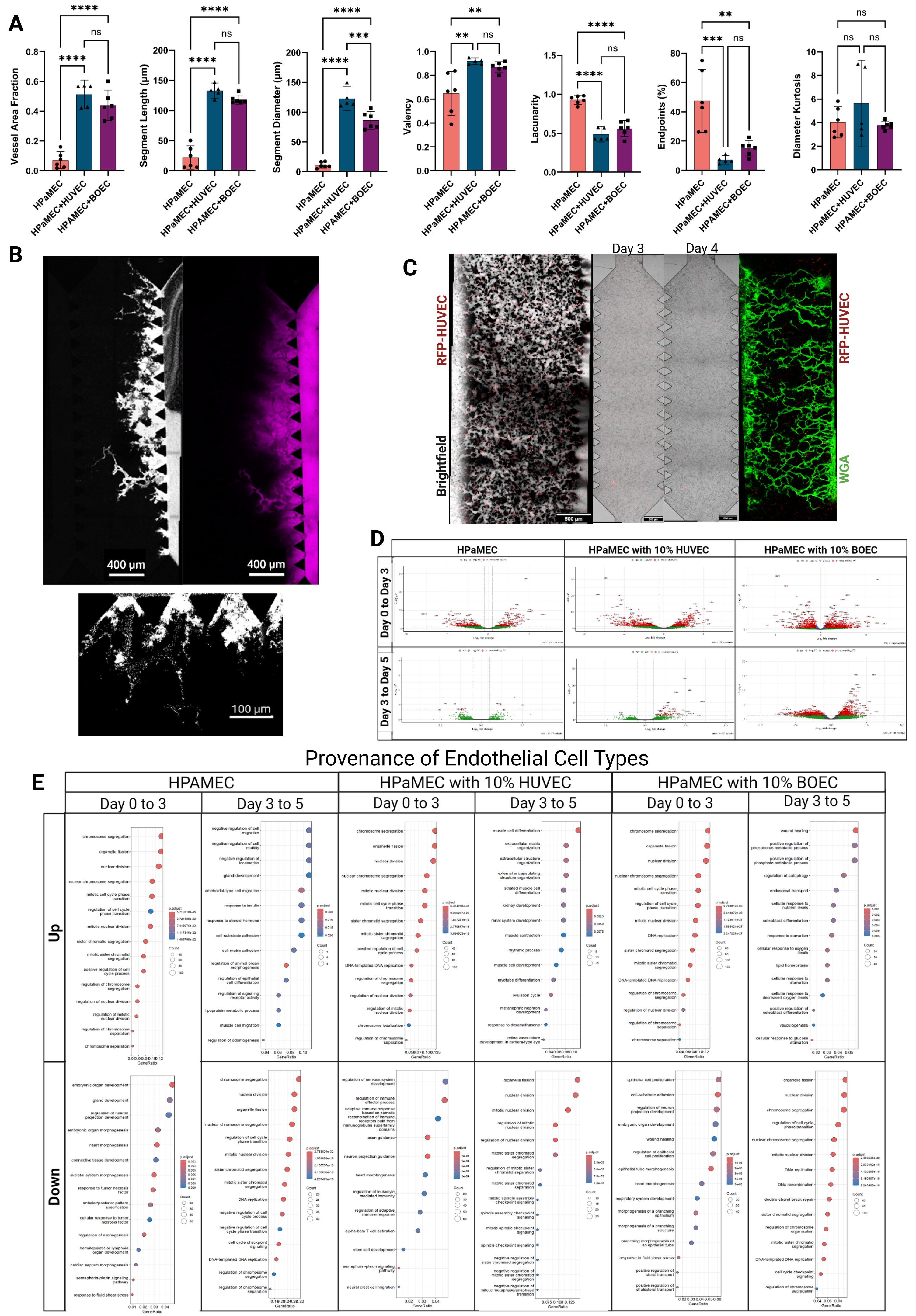

### Supplementary Figure 4

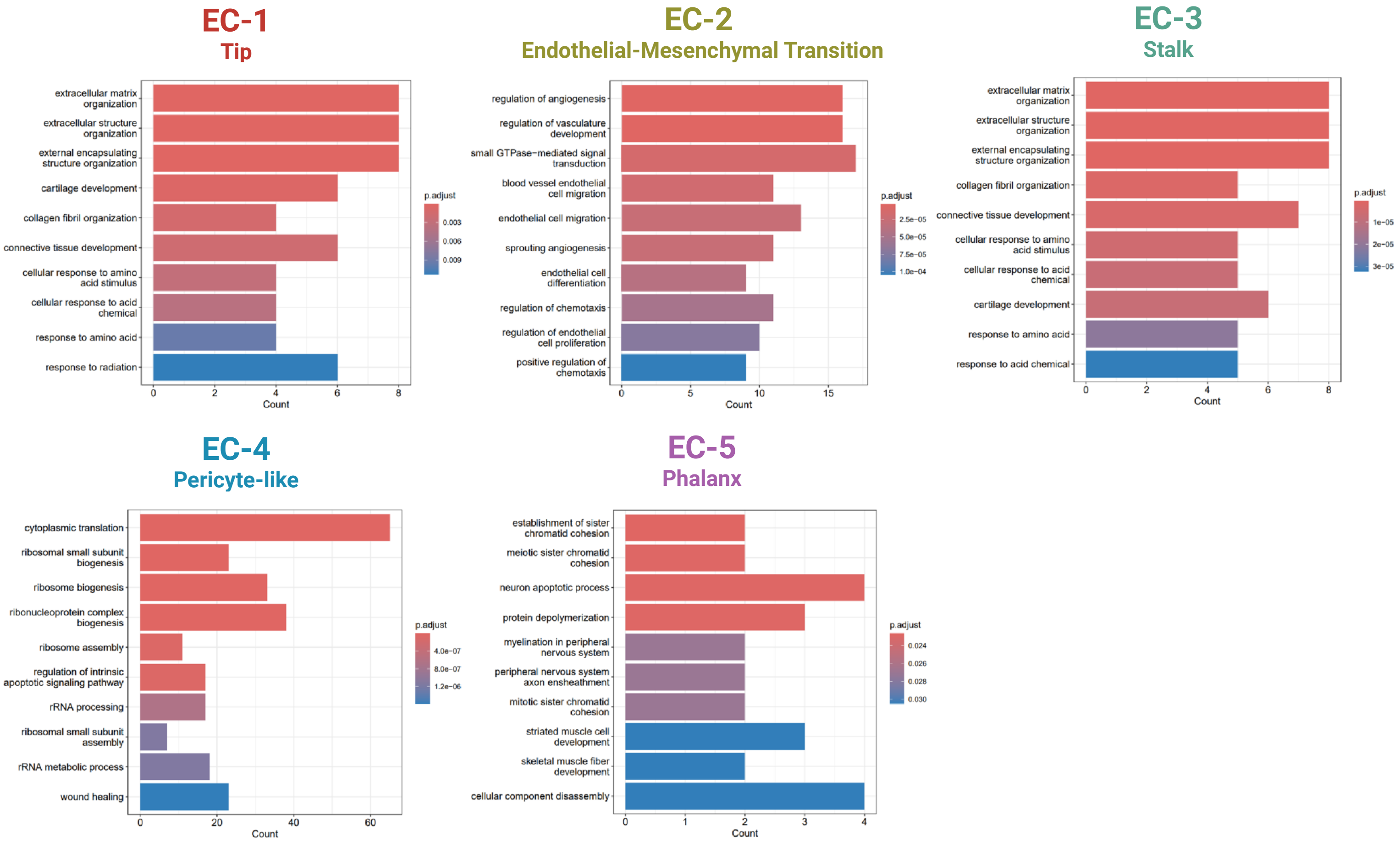

### Supplementary Figure 5

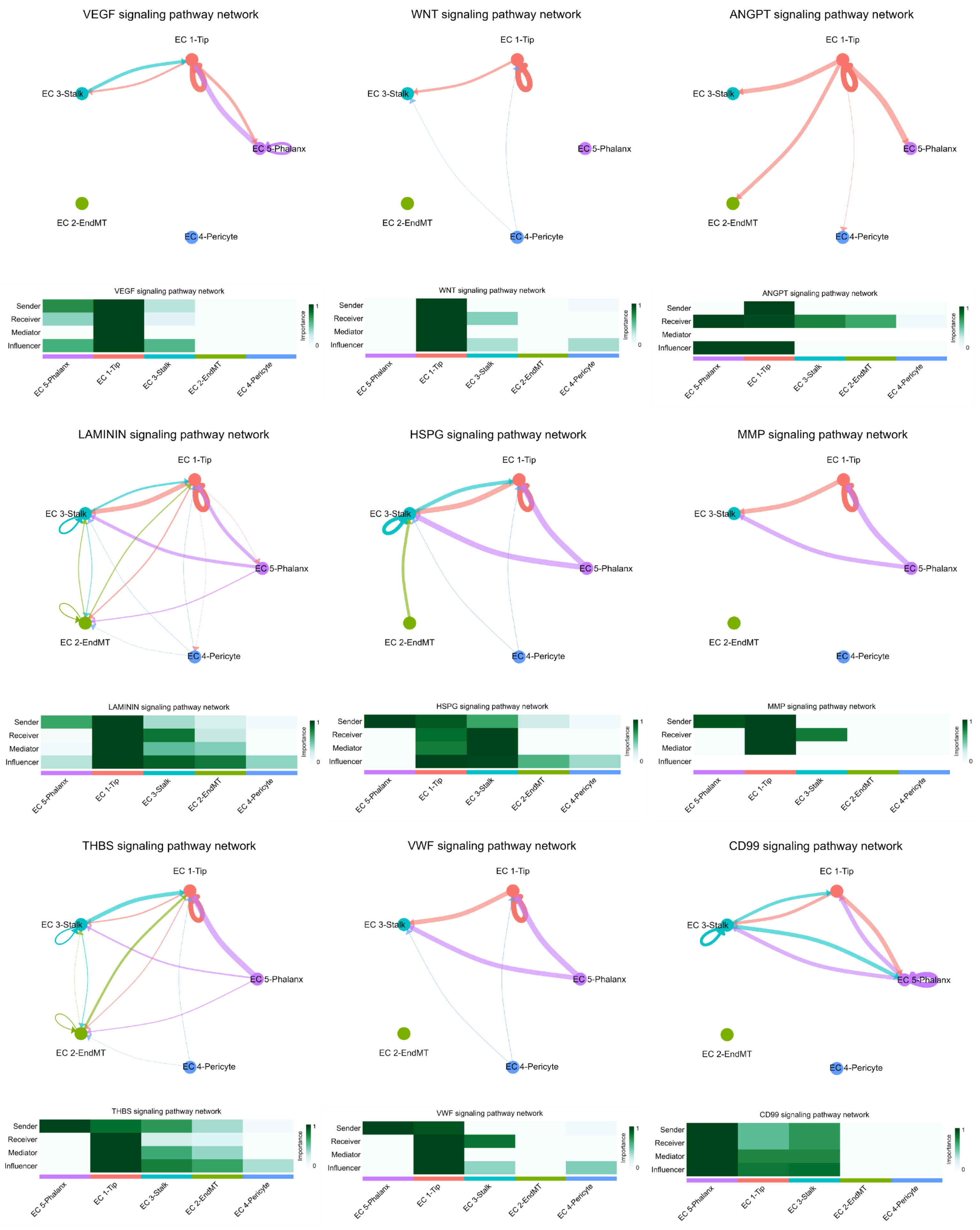

### Supplementary Figure 6

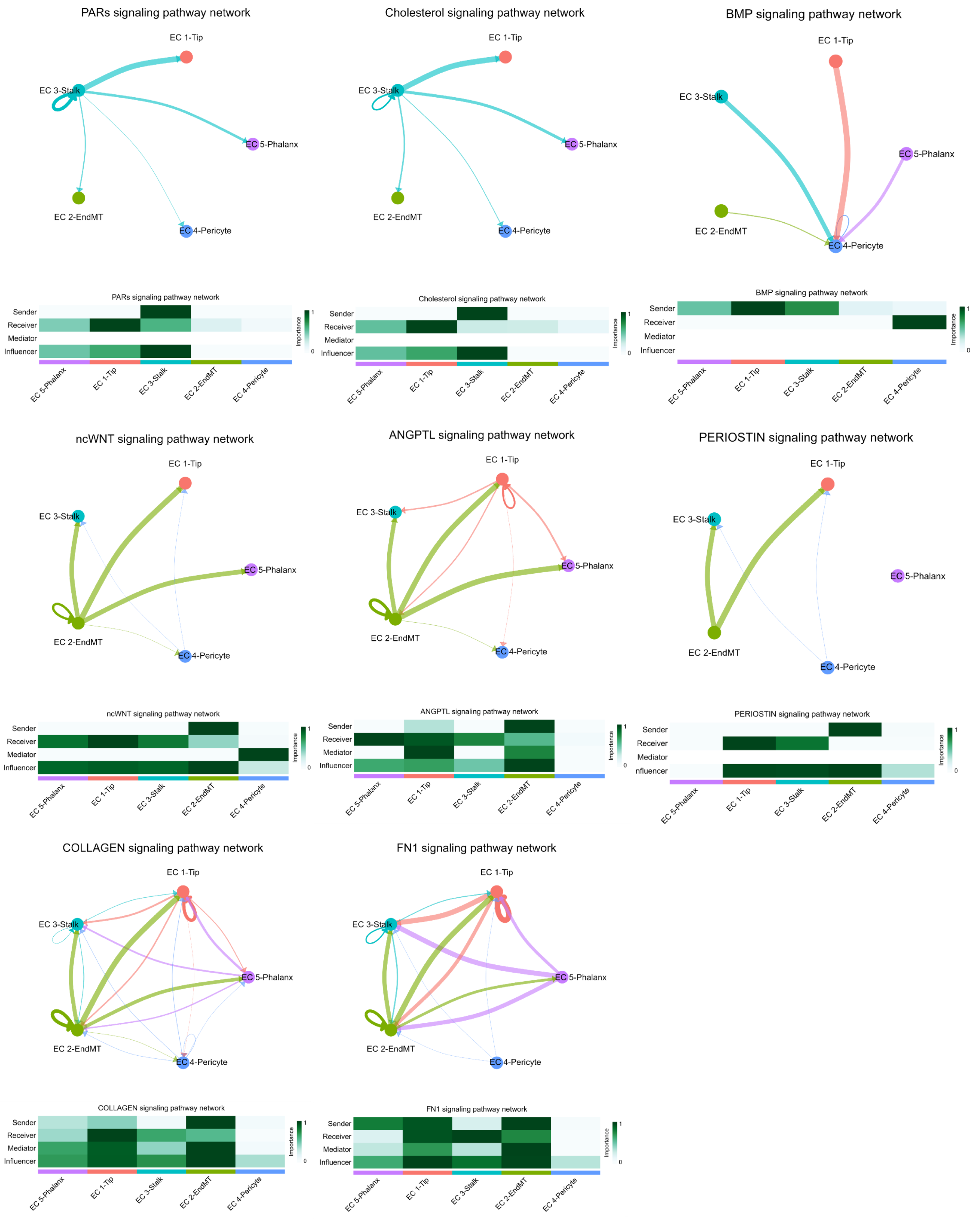

### Supplementary Figure 7

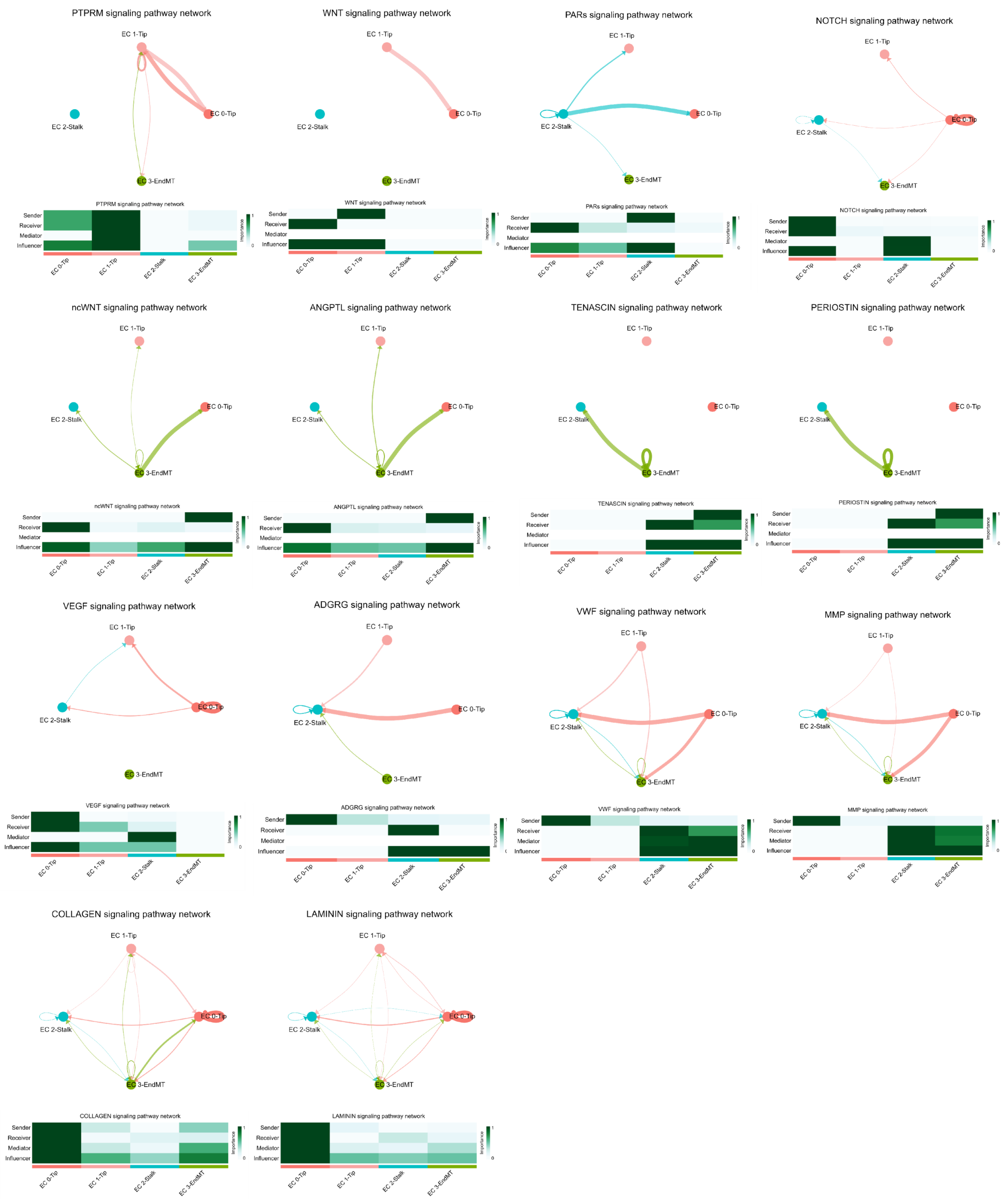

### Supplementary Figure 8

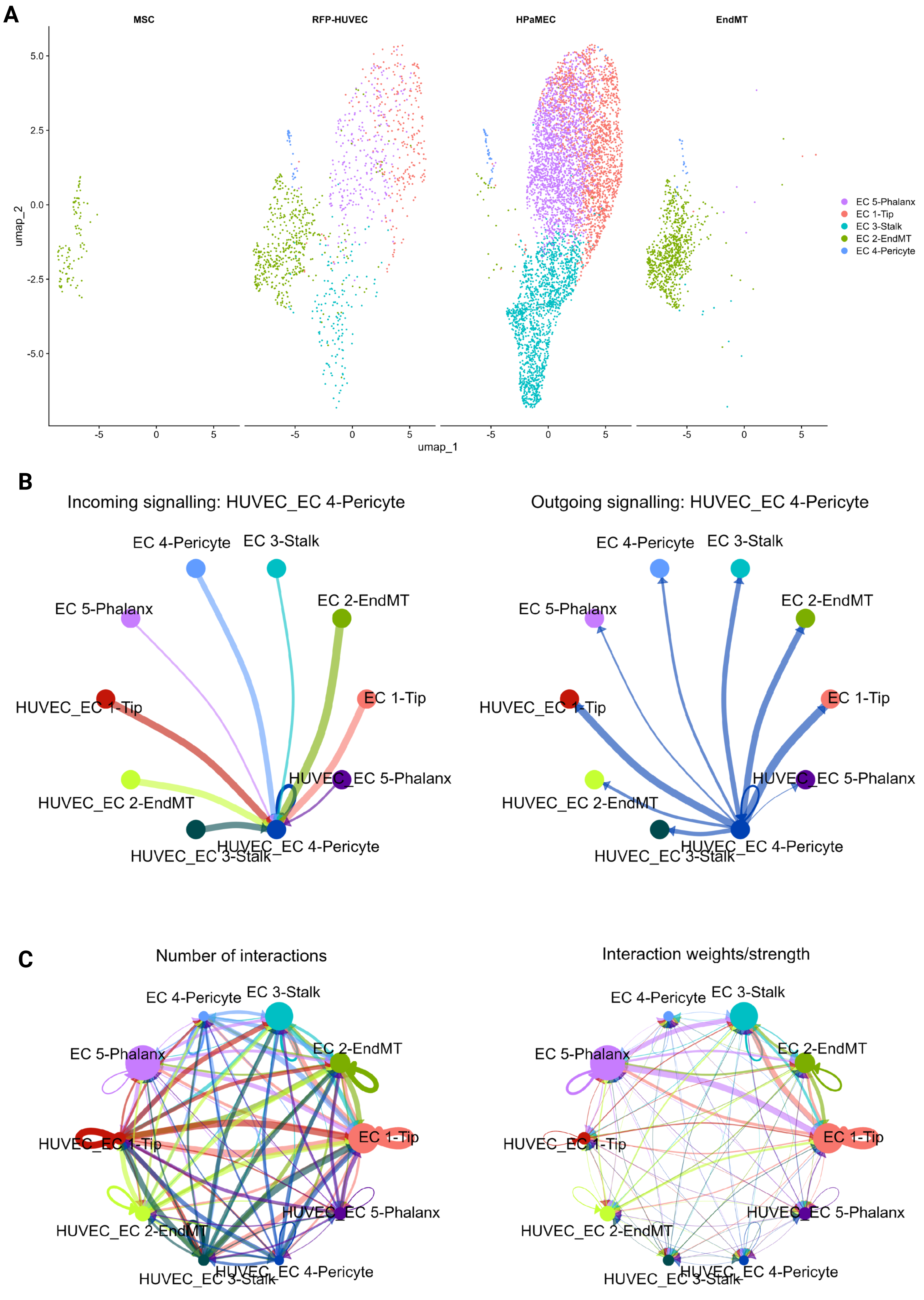
